# Forsythoside A blocks the infection of Avian Infectious Bronchitis Virus through binding S1 subunit

**DOI:** 10.1101/2023.09.15.557895

**Authors:** Ruiting Shen, Xuewei Liu, Peng Yin, Xu Li, Jun Chu, Hua Yao, Xiaolin Hou

**Author notes:** Corresponding author: Xiaolin Hou. Ruiting Shen, Xuewei Liu, Peng Yin, and Xu Li contributed equally to this work.

## Abstract

Avian infectious bronchitis (IB) is an acute highly contagious respiratory infectious disease caused by avian infectious bronchitis virus (IBV), causing serious economic losses to poultry industry. Forsythiaside A (FTA) is a phenylethanol glycoside compound in the fruit of *Forsythia suspensa*, which is widely used for inflammatory respiratory tract disease, including viral infected inflammation. In this study, the mechanism of FTA against IBV was researched. To determine the inhibitory effect of FTA on IBV infection and the interaction between FTA and IBV particles, immunofluorescence assay, plaque assay, RT-qPCR and enzyme-linked immunosorbent assay (ELISA) were performed and it was discovered that FTA inhibited IBV infection and viral attachment to host receptor under non-cytotoxic concentrations, and FTA may bind to virus particles. Then ELISA and isothermal titration calorimetry (ITC) were employed to further study the interaction between FTA and S protein. The results indicated that FTA bound to S1 subunit rather S2 subunit. Besides, the molecular docking showed that FTA form hydrogen bonds with eight amino acids in the S1 subunit. When these amino acids were mutated. the affinity disappeared between the S1 mutant and FTA. In conclusion, FTA prevents IBV infection by blocking viral attachment and contacts with the S1, but not the S2 subunit of IBV. At last, the bind sites were identified by comparing the affinity of wild S1 and mutant S1.

## 1. Introduction

Avian infectious bronchitis (IB) is an acute highly contagious respiratory infectious disease caused by avian infectious bronchitis virus (IBV)^[1]^. IB not only lowers egg production and quality but also has a negative impact on the worldwide economy, making it a serious issue for the poultry industry^[2]^. The most common symptoms of IB are related to the respiratory tract, including wheezing, coughing, sneezing, trachea rale and runny nose^[3]^. Although the main target of IBV is the epithelial surface of the respiratory tract, it can replicate in other organs, including the kidney, gastrointestinal tract, urinary tract and reproductive tract, depending on the virus strain^[4]^.

At present, vaccination is still the most popular way to prevent production loss. However, due to the poor cross protection to mutable and different serotype IBV strains, the current vaccines are insufficient to provide adequate protection^[5]^ Forsythiaside A (FTA) is the main active ingredient isolated from *Forsythia suspensa*, which has significant biological activity, showed notable pharmacological properties in inflammatory disease, virus infection, neurodegeneration, oxidative stress, liver injury, and bacterial infection^[6]^ In 2019, coronavirus disease 2019 (COVID-19) pneumonia began widly prevalent by SARS-CoV-2. FTA as a part of Lianhuaqingwen capsule, a herb medicine product, was found that angiotensin-converting enzyme 2 (ACE2), the receptor of SARS-CoV-2, could be inhibited by the component. Besides, the molecular docking result showed that FTA could bind to the active region of ACE2 which interact with viral S protein (Asp350 to Arg393)^[7]^. The main protease (Mpro) of SARS-CoV-2 is an important non-structural protein mediates viral replication and transcription. It was found that the Mpro enzymatic activity was inhibited at IC_50_=22.26 μM of FTA and the docking results indicated that FTA was able to bind some sites of Mpro. What’s more, FTA also had a direct virucidal effect on IBV^[9]^. But at the detailed anti-IBV role and the potential target remain unknown.

## 2. Materials and Methods

### 2.1. Cells, viruses, and reagents

African green monkey kidney cells (Vero) and Human Embryonic Kidney 293T (HEK293T) were cultured in Dulbecco’s Modified Eagle Medium (DMEM, Gibco, USA) supplemented with 10% Fetal Bovine Serum (FBS, HyClone, USA) containing penicillin (100 units/ml) and streptomycin (100 μg/ml) (Gibco, USA). All cells were grown at 37 °C containing 5% CO_2_ in incubator.

The IBV strain Beaudette (GenBank accession number: MZ368698.1), which is maintained in our laboratory, was used for all experiments and is represented as “IBV” in this article.

Forsythiaside A (FTA), purity ≥ 98%, was used for the in this study (Desite, China).

### 2.2. Virus titration

Vero cells grown in 96-well plates were infected with ten-fold serial dilutions of IBV. After 1 h at 37 °C, the culture medium was replaced with fresh DMEM-2% FBS. Viral titers were determined using endpoint dilution analysis at 5 days post-inoculation. The Reed–Muench method was used to calculate the 50% tissue culture infected dose (TCID_50_).

### 2.3. Cytotoxicity assay

The cytotoxic effects of FTA on Vero cells were measured by MTT assay. Briefly, Vero cells were seeded and grown to 90% confluence in 96-well cell culture plates (Corning Incorporated, USA), the supernatant was discarded and the cells were washed with Phosphate Buffered Saline (PBS, HyClone, USA) three times. Different concentrations of FTA were incubated with Vero cells at 37 °C for 48 h. Then the medium was discarded, 100 μL of MTT (1 mg/mL) (Solarbio, China) was added in each well and maintained at 37 °C for 4 h. The formed formazan crystals were dissolved in 150 μL dimethyl sulfoxide (DMSO) (Solarbio, China). The OD value was measured with a microplate reader at 490 nm. The percentage of surviving cells were calculated and used the following formula: cell viability (%) = (OD_490nm_ drug) / (OD_490nm_ control) × 100%. The highest nontoxic concentration of FTA was chosen as the most safety concentration. The experiments were performed in triplicate.

### 2.4. Quantitative reverse transcription PCR (RT-qPCR)

Total RNA was extracted from IBV-infected cells at 8 hour-post-infection (h.p.i.), using FastPure Cell/Tissue Total RNA Isolation Kit V2 (Vazyme, China). Subsequently, Evo M-MLV RT Premix for qPCR (Agbio, China) was used to synthesize cDNA and quantification of viral gene expression was performed on the Mx3005P Real-Time PCR System (Stratagene, USA), using Ultra SYBR Mixture (Low ROX) (CWBIO, China). The genomic of IBV Beaudette Nucleocapsid was detected by the following primers: forward: 5’-GAACAGGACCAGCCGCTAA-3’ and reverse 5’-GAGGAATGAAATCCCAACG-3’. β-actin gene was used as a reference gene and detected using the primers: forward: 5’-AGGCATCCTCACCCTGAAGTACC-3’ and reverse 5’-CCACACGCAGCTCATTGTAGAAGG-3’. The qPCR reaction procedure as follows: 10 min at 95 °C, 40 cycles of 15 s at 95 °C and 60 s at 60 °C, then 15 s at 95 °C, 60 s at 60 °C and 15 s at 95 °C. Each sample was performed in triplicate. The relative quantitation of IBV-N gene expression was determined by comparison to β-actin.

### 2.5. Virus pre-treatment assay

To observe the effect of FTA treatment at different times, plaque assay and RT-qPCR were performed. With the intent to assess the preventive effect of FTA, Vero cells were seeded in 6-well plates and treated with concentration of 50 μM, 100 μM or 200 μM of FTA for 12 h at 37 °C respectively, then the drug was abandoned, and cells were washed three times with PBS. 100TCID_50_ of IBV was incubated with Vero at 37 °C for 1 h, then removed the virus and washed three times. For plaque assay, cells were overlaid with DMEM containing 2% methylcellulose (Solarbio, China) and 2% FBS and incubated for 48 h at 37 °C. The cells were overlaid with 1% crystal violet (Solarbio, China) and incubated for an additional 30 min at room temperature. Plaque-forming units were counted. The viral infection rate was calculated as the formula: Viral infection rate (%) = (the number of plaques of FTA treated)/ (the number of plaques of IBV infected) × 100%. DMEM treatment was included as control in each condition. All the experiments were performed in triplicate.

### 2.6. Virus attachment assay

With a view of study whether FTA could inhibit attachment of IBV, Vero cells were prechilled at 4 °C for 1 h and subsequently challenged with 100TCID_50_ of IBV in the presence of FTA for 1 h at 4 °C. After washed three times with ice-cold PBS, the cells were coated with 2% DMEM at 37 °C for 48 h. Viral plaque assay was performed. The viral infection rate was calculated as the formula: Viral infection rate (%) = (the number of plaques of FTA treated)/ (the number of plaques of IBV infected) × 100%. DMEM treatment was included as control in each condition. All the experiments were performed in triplicate.

### 2.7. Virus entry assay

To evaluate the effect of FTA on the entry of IBV, Vero cells were prechilled at 4°C for 1 h before inoculation with 100TCID_50_ of IBV for 1 h at 4 °C. The cells were then treated with FTA and further incubated at 37 °C for 1 h. At the end of the incubation, extracellular virus was inactivated by citrate buffer (pH 3.0) for 1 min and then washed with PBS three times before overlaying with 2% DMEM for 48 h at 37 °C. For plaque assay, The viral infection rate was calculated as the formula: Viral infection rate (%) = (the number of plaques of FTA treated)/ (the number of plaques of IBV infected) × 100%. DMEM treatment was included as control in each condition. All the experiments were performed in triplicate.

### 2.8. Virus post-infection assay

To investigate the effect of FTA after viral entry, Vero cells were infected with 100TCID_50_ of IBV for 1h at 37°C, then washed before overlaid medium containing 50 μM, 100 μM, 200 μM of FTA. Viral plaque assay was performed. The viral infection rate was calculated as the formula: Viral infection rate (%) = (the number of plaques of FTA treated)/ (the number of plaques of IBV infected) × 100%. DMEM treatment was included as control in each condition. All the experiments were performed in triplicate.

### 2.9. Viral inactivation assay

100TCID_50_ of IBV was mixed with 50 μL of 50 μM, 100 μM, 200 μM of FTA respectively and incubated at 37 °C for 1 h. The mixture was then diluted 20-fold with no phenol red DMEM, and the dilution was subsequently added to the monolayers of Vero cells which were seeded in 6-well plates. After adsorption for 1 h at 37 °C, the medium was discarded, and the cells were washed with PBS three times. Then the cells were overlaid with the maintenance medium and further incubated at 37 °C for 48 h. Viral plaques were counted as previously described. The experiments were performed in triplicate.

### 2.10. Immunofluorescence assay (IFA)

Vero cells were seeded in 12-well plates including 20 mm coverslips (Solarbio, China) and pre-treated with 50 μM, 100 μM or 200 μM of FTA, respectively, for 1 h at 37 °C and then infected with IBV for 1 h at 37 °C. Cells were washed with PBS and then incubated in fresh medium containing FTA for 48 h. DMEM served as the treatment control. Subsequently, the overlay was removed, and the coverslips were rinsed three times with PBS. The cells were then fixed with 4% PFA (Solarbio, China) for 15 min at room temperature (RT) and blocked with QuickBlock™ Blocking Buffer (Beyotime, China) for 1 h. The anti-IBV-chicken positive serum (B0116, QYH Biotech, China) was used as primary antibody (1:500) and incubated at 4°C overnight. Rabbit anti-chicken IgG conjugated with Cy3 (K1035R-Cy3, Solarbio, China) was used as the secondary antibody (1:1000) and incubated at room temperature for 1 h in the dark. Hoechst 33342 (C1022, Beyotime, China) was used to stain the cell nuclei for 5 min at room temperature. Coverslips were mounted on a microscope slide and viewed with Leica Confocal Microscopes Stellaris 5. Representative images were obtained and processed using the LAS X Navigator software.

### 2.11. Alignment of FTA and IBV S protein

The 3D structural diagram of IBV S protein (6CV0) was taken from Protein Data Bank (http://www.rcsb.org/pdb/), and the 3D structure diagram of FTA was drawn by Chem3D software. Autodock was used to calculate and the docking result was showed by PyMOL.

### 2.12. Plasmid construction

Genes encoding IBV-Beaudette-S1 (the aa 1-532 of precursor of S protein) and IBV-Beaudette-S2 (the aa 539-1091 of precursor of S protein) were amplified from Vero cells infected with IBV-Beaudette by reverse-transcription PCR (RT-PCR). The amplified fragment was ligated into the pCDNA3.4 vector (Addgene, USA) with a C-terminal 6×His-tag tag. The IBV-Beaudette-S1 mutant [(aa 434, 440, 441, 454, 469, 471, 473, 485 were mutated to phenylalanine (aa 434), alanine (aa 440), alanine (aa 441), valine (aa 454, aa 469 and aa 471), asparagine (aa 473) and phenylalanine (aa 485)] was synthesized by Detai Bioscience, Inc.

### 2.13. Expression and purification of recombinant proteins

The pCDNA3.4 expression vectors containing the described sequences above was transfected into HEK293T cells by the measure of GP-transfect-Mate (GenePharma, China) for 48 h at 37 °C. Then the medium was discarded, and the cell lysate was harvested. The lysate of HEK293T with no plasmid was regarded as control. The proteins were purified by the manual of Ni-IDA agarose gel (Detai Bioscience, China), and the products were stored at −80 °C.

### 2.14. SDS-PAGE and Western blotting

The expression and purification of the proteins were analyzed by Western blotting and sodium dodecyl sulfate-polyacrylamide gel electrophoresis (SDS-PAGE). The cell lysate was mixed with 5 × SDS-PAGE Sample Loading Buffer (Beyotime, China) and boiled at 100 °C for 10 min. Equal amounts of protein were separated on a 10% SDS polyacrylamide gel and electro-transferred from the gel to a nitrocellulose filter (NC) membrane. After blocking with 5% skimmed milk powder in Tris-HCl buffer with Tween-20 (TBST) (LABLEAD, China), the membrane was incubated with Rabbit His-Tag antibody (1:1000) (#12698, Cell Signaling Technology, USA) at 4 °C overnight and IRDye® 800CW Goat anti-Rabbit IgG (1:15000) (926-32211, LI-COR, USA) at room temperature for 1 h. β-actin was used as the internal loading control. The membrane was scanned by Odyssey Infrared Imaging (LI-COR, USA). The purified protein was mixed with Loading Buffer and boiled. 10% SDS polyacrylamide gel was used to separate the protein and then stained by Coomassie blue solution for 30 min and the gel was decolorized by ddH_2_O overnight.

### 2.15. The conjugation of FTA and OVA

FTA (0.04 mmol), succinic anhydride (0.04 mmol) (Leyan, China) and triethylamine (0.04 mmol) (Sinopharm, China) were dissolved in N, N-Dimethylformamide (DMF) (500 μL) (Macklin, China) and pyridine (500 μL) (Macklin, China). The mixture was stirred at room temperature for 2 h, then the mixture was evaporated by vacuum drying oven at room temperature and the crystals were collected.

The crystals were completely dissolved in 500 μL DMF, then 0.2 mmol 1-(3-Dimethylaminopropyl)-3-ethylcarbodiimide hydrochloride (EDC) (Sigma, USA) and 0.2 mmol N-Hydroxy succinimide (NHS) (Macklin, China) were added while stirring, and the mixture was stirred at room temperature overnight. The supernatant was collected after centrifugation which was regarded as solution A. 15 mg ovalbumin (OVA) (Macklin, China) was dissolved in 5 mL PBS which was regarded as solution B. Under magnetic stirring, solution A was added drop by drop to solution B and mixed at room temperature for 12 h. When the reaction finished, the supernatant was collected and dialyzed at 4 °C for 3 d by dialysis bag (14000 kD, Beyotime, China), changed dialysate (PBS) three times a day. The solution in the dialysis bag was obtained and the precipitate was discarded. To determine whether the conjugation of FTA and OVA was successful, the 190 nm - 400 nm spectrum of the mixture was scanned by DeNovix DS-11 Spectrophotometer/Fluorometer Series.

### 2.16. ELISA

FTA-OVA in carbonate buffer (15 mM Na_2_CO_3_, 35 mM NaHCO_3_, pH 9.6) were coated in 96 well flat bottom polystyrene plates and incubated at 4 °C overnight. Then the plates were washed three times with 200 μL Phosphate Buffered Saline with Tween-20 (PBST) and blocked for 1 h at room temperature with 100 μL 5% skimmed milk powder in PBS. After that the plates were washed three times with PBST and two-fold diluted IBV, IBV-Beaudette-S1 or IBV-Beaudette-S2 were added in plate and incubated for 1 h at 37 °C. PBS or HEK293T lysis were used as negative control. Followed three times washed with PBST, primary antibody [Infectious Bronchitis virus Nucleoprotein antibody (1:1000) (B819M, GeneTex, USA) or Rabbit His-Tag antibody (1:1000) (#12698, Cell Signaling Technology, USA)] and secondary antibody [Goat anti-mouse IgG conjugated with HRP (1:1000) (SE131, Solarbio, China) or Goat anti-rabbit IgG conjugated with HRP (1:1000) (SE134, Solarbio, China)] were incubated for 1 h at 37 °C, respectively. 100 μL TMB (Solarbio, China) was added in each well at room temperature for 60 min and the reaction was terminated by Stop Solution for TMB Substrate (Solarbio, China). The absorbance was immediately determined at 450 nm. The OD_450nm_ was calculated and the P/N=2.5-3.0 was regarded as positive.

### 2.17. Isothermal titration calorimetry (ITC)

FTA was prepared as 1 mM in PBS and loaded in sampling needle. S1 protein (1 μM) was dissolved in PBS then added to the sample cell. The titration reaction temperature was 25 °C, the number of drops was set to 20, reference power was 10 μcal/s, single titration volume was 2 μL, the single injection time of FTA was 4 s, the interval time between two titrations was 180 s, and the agitator speed was 1000 rpm/min. When titrating S2 protein, the concentration of S2 protein was 1 μM. FTA concentration was changed into 100 μM, the titration interval was set to 60 s, and other conditions remain unchanged.

### 2.18. Statistical analysis

All data were expressed as the mean ± standard error of the mean (SEM) from three independent experiments performed in triplicate. The statistical analyses were conducted using Student’s t-test in GraphPad Prism version 8.0 (GraphPad Software, San Diego, CA, USA). A p-value was considered significant as the following: \**p* < 0.05; \*\**p* < 0.01; \*\*\**p* < 0.001; \*\*\*\**p* < 0.0001)

## 3. Results

### 3.1. The cytotoxic effect of FTA on Vero cells

To investigate whether FTA treatment affects cell viability, the toxicity of FTA on Vero cells was determined by MTT assay. As the data shown, the cell viability was slightly decreased at 300 μM, and the viability was decreased approximately 20% and 40% at the concentration of 400 μM and 500 μM, respectively (Fig.1). These results indicated that FTA did not influence cell viability at concentrations below 300 μM. Therefore, the concentration of 50 μM, 100 μM and 200 μM of FTA were chosen for the subsequent antiviral assays.

**Fig.1.**
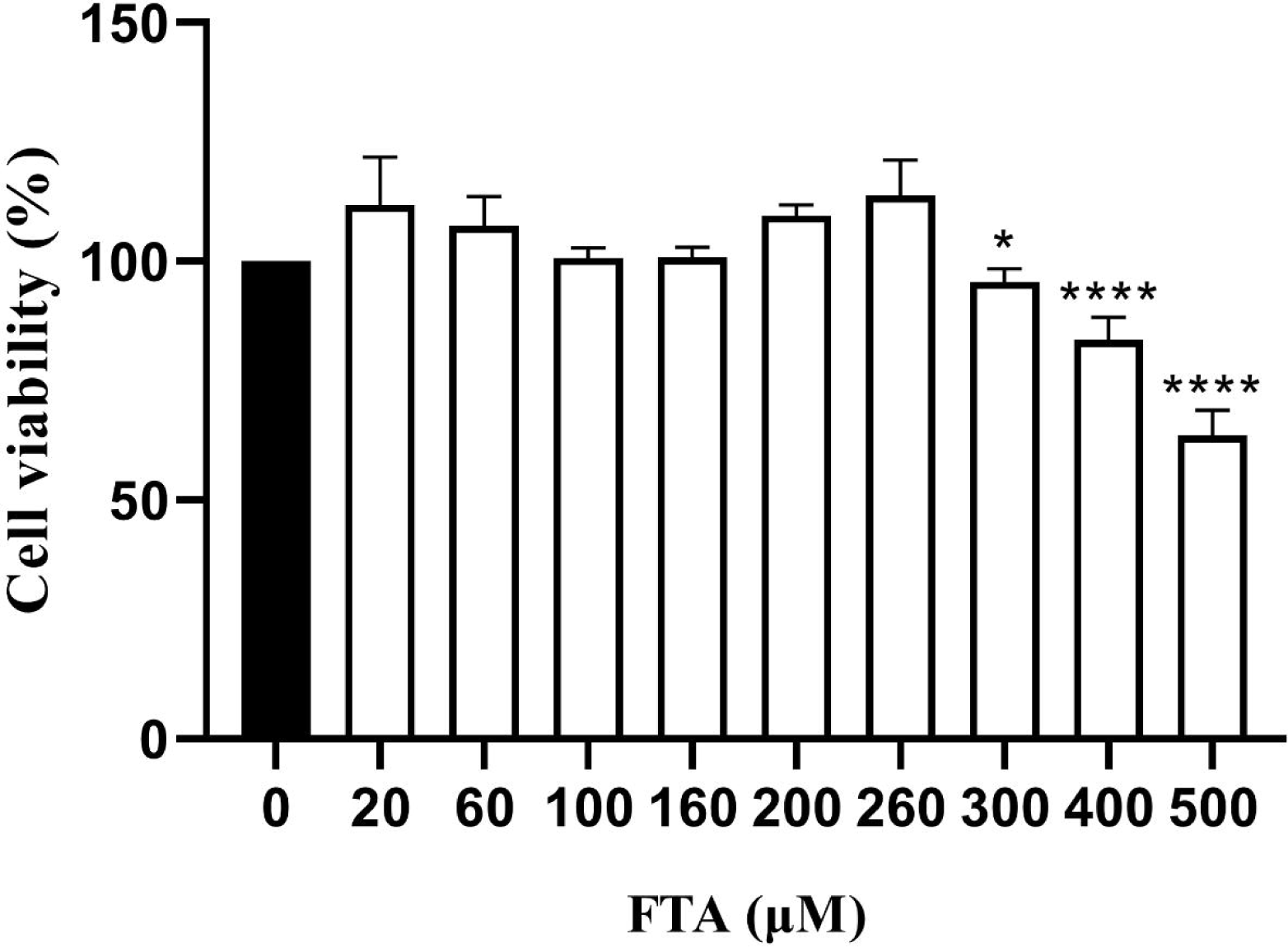
FTA had cytotoxicity on Vero cells. Vero cells were treated with different concentrations of FTA and cell viability was detected by MTT assay. The experiment was repeated three times. \**p* < 0.05, \*\*\*\**p* < 0.0001, compared with 0 μM.

### 3.2. FTA inhibits IBV infection

Immunofluorescence assay (IFA) was performed to study whether FTA could inhibit IBV infection. As the result shown, when the concentration of FTA was increased, the red fluorescence,hich indicated as IBV particles, was faded (Fig.2), indicating that FTA has an effect of anti-IBV infection, but the mechanism not revealed.

**Fig.2.**
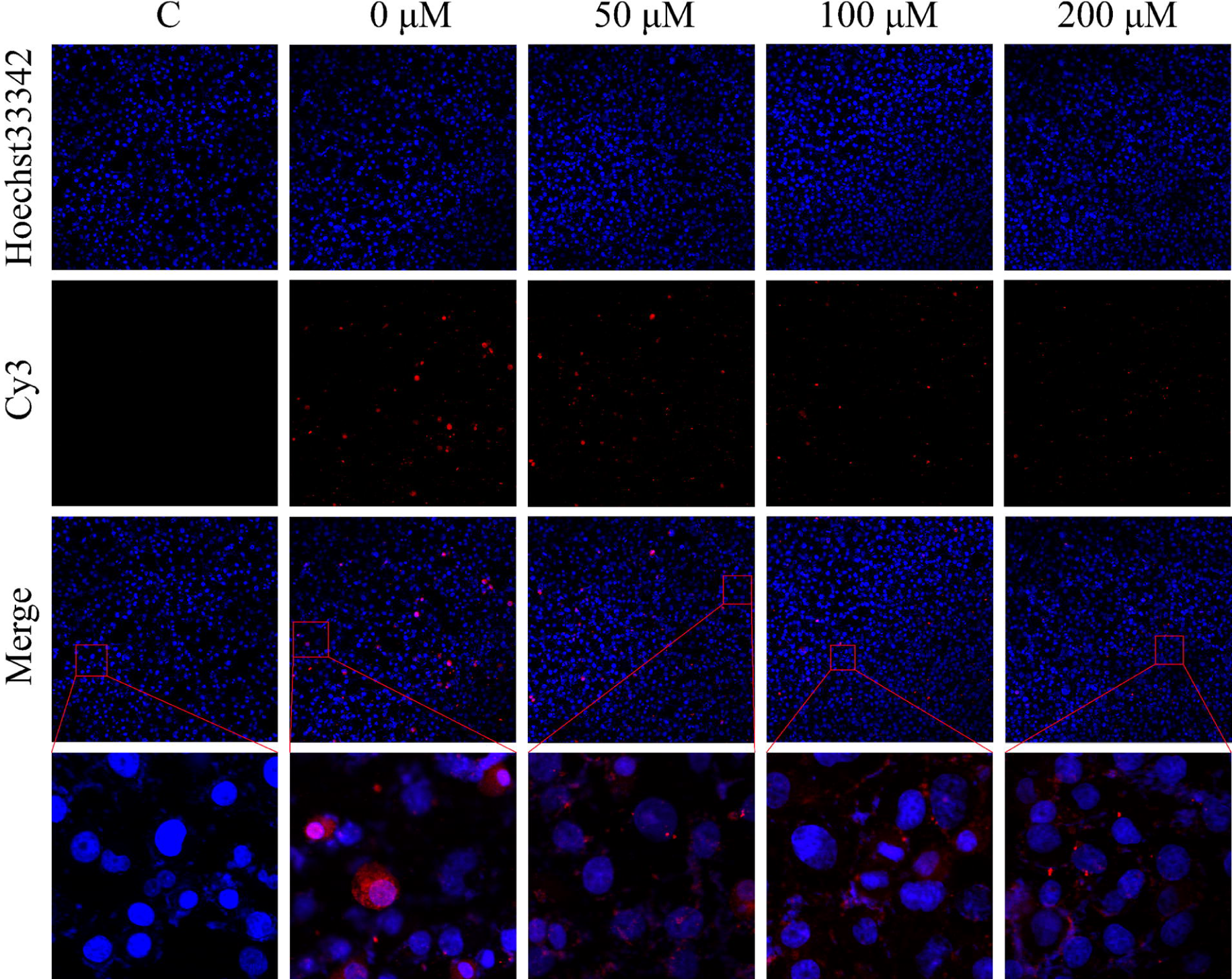
Identification of antiDIBV activity of FTA in Vero cells. Vero cells were pre-treated with the indicated concentrations of FTA for 1 h and then infected with IBV for 1 h at 37 °C. Cells were washed with PBS and then incubated in fresh medium containing FTA for 48 h. DMEM served as the treatment control. At 48 h.p.i., IFA images of Vero IBV-infected and FTA-treated cells were collected. Cell nuclei were indicated by Hoechst33342 for blue, IBV particles were indicated by Cy3 for red.

### 3.3. FTA inhibits the attachment of IBV and may bind to IBV particles

To better understand the antiviral mechanism and the stage of IBV infection affected by FTA *in vitro*, plaque assay and RT-qPCR were performed. As the concentration of FTA increased, the plaque formation of stage of viral attachment decreased. However, the plaque formation of the stages of FTA pretreatment, viral entry and viral post-infection were not decreased by FTA treatment (Fig.3). After plaques were counted and RT-qPCR was performed, the results showed that there was no inhibition effect of FTA on IBV entry (Fig.4C, Fig.4H), not prevent IBV infection either (Fig.4A, Fig.4F) and have no effect in the post-infection stage. (Fig.4D, Fig.4I). But viral attachment was decreased by FTA treatment (Fig.4B, Fig.4G).

**Fig.3.**
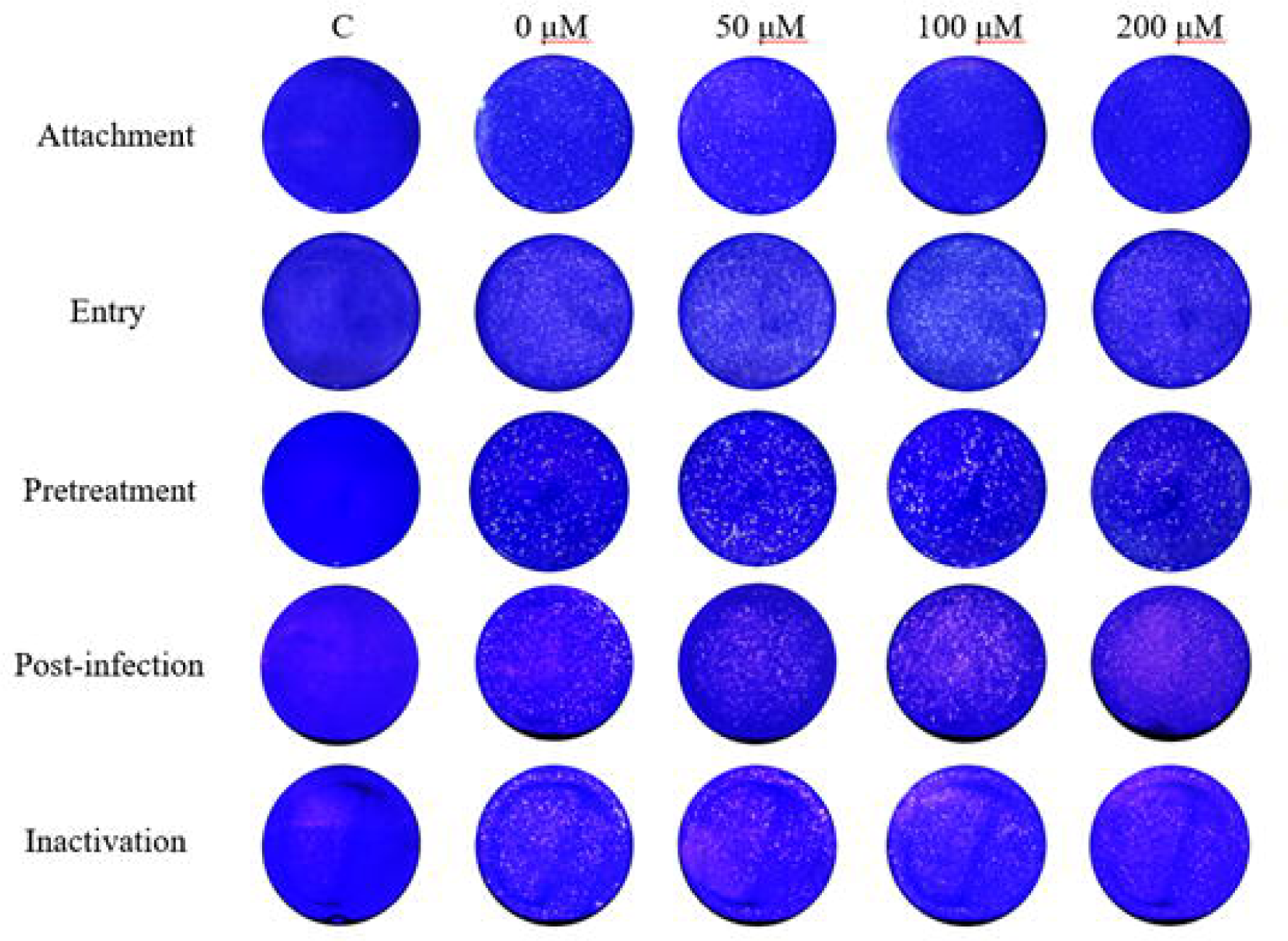
The effect of FTA addition time on the plaque formations. Plaque formations of plaque assay of each stage. The experiment was repeated three times.

**Fig.4.**
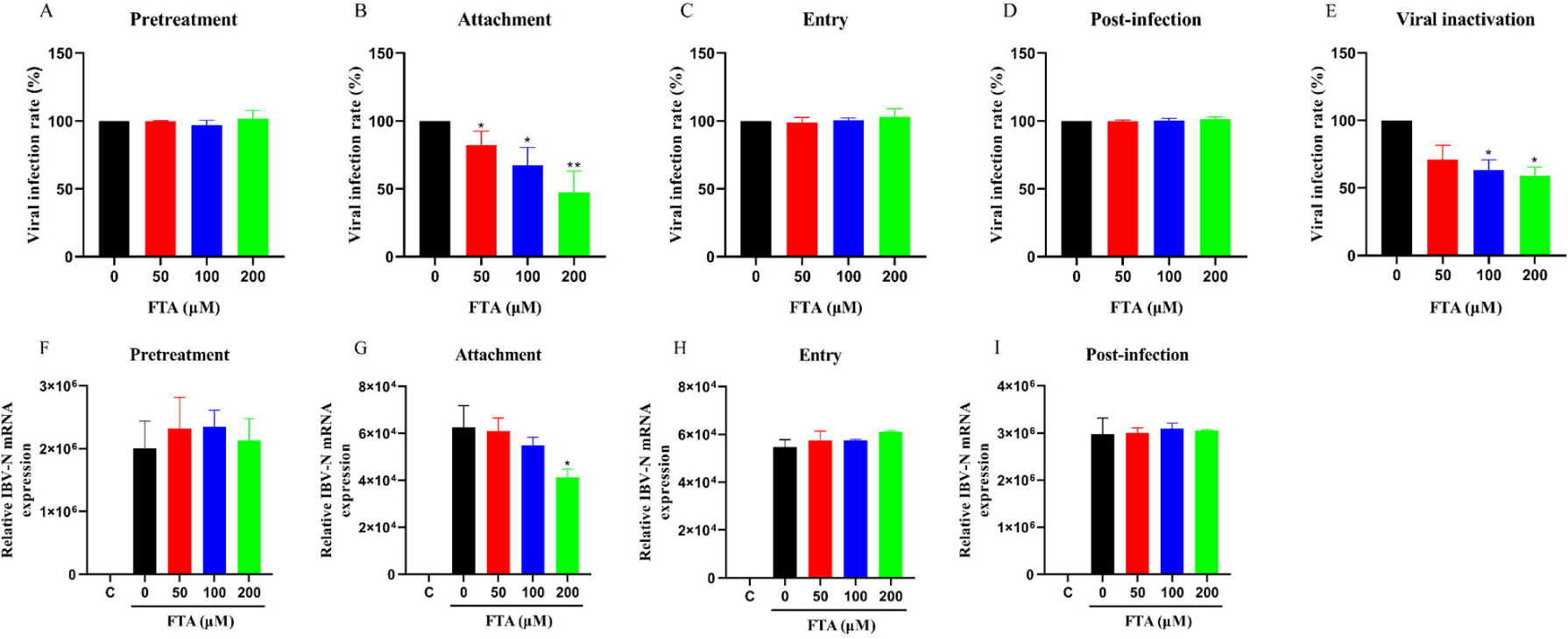
The effect of FTA on the different stages of IBV infection. Plaque assay and RT-qPCR were used to determine the stage of IBV infection. (A, F) FTA pre-treatment assay. (B, G) Viral attachment assay. (C, H) Viral entry assay. (D, I) Viral post-infection assay. (E) Viral inactivation assay. The experiment was repeated three times. \**p* < 0.05, \*\**p* < 0.01, compared with 0 μM.

The adsorption of IBV requires interaction between S protein and receptors (alpha2,3-linked sialic acids) on the host cell^[10]^. However, the pretreatment by FTA for 12 h didn’t reduce the infection of IBV on Vero cells. This result indicated that there was not an interaction between FTA and cell, so we assumed that the efficacy of FTA can only be achieved by combining with IBV particles. To verify this hypothesis, different concentrations of FTA were mixed with 100 TCID_50_ IBV at 37 °C for 1 h, then the mixtures were diluted 20-fold to avoid the effect to cells. The plaque assay result showed that viral infection rate decreased when the concentration of FTA increased (Fig.3, Fig.4E). The results indicated that the inactivation of IBV infection by FTA is characterized by reduced attachment, and FTA may bind to IBV particles.

### 3.4. Synthesis of artificial antigen FTA-OVA

Hapten refers to a low molecular weight compound that cannot trigger the immunogenic reaction of producing antibodies and should be covalently coupled with immunogenic macromolecules (carrier protein) to obtain immunogenicity. According to this, a carboxyl needs to be introduced into FTA using the succinic anhydride method, to further synthesize artificial antigens. After the hapten was contributed and conjugated with OVA, FTA-OVA was determined by a spectrophotometer. Red indicated the spectrum of OVA, with a characteristic absorption peak at approximately 228 nm; Blue represented the spectrum of FTA, with a characteristic absorption peak at approximately 327 nm; Green was the spectrum of the synthesized artificial antigen. It has two absorption peaks at about 203 nm and 349 nm, including the characteristic absorption peaks of OVA and FTA, indicated that the artificial antigen was successfully synthesized (Fig.5).

**Fig.5.**
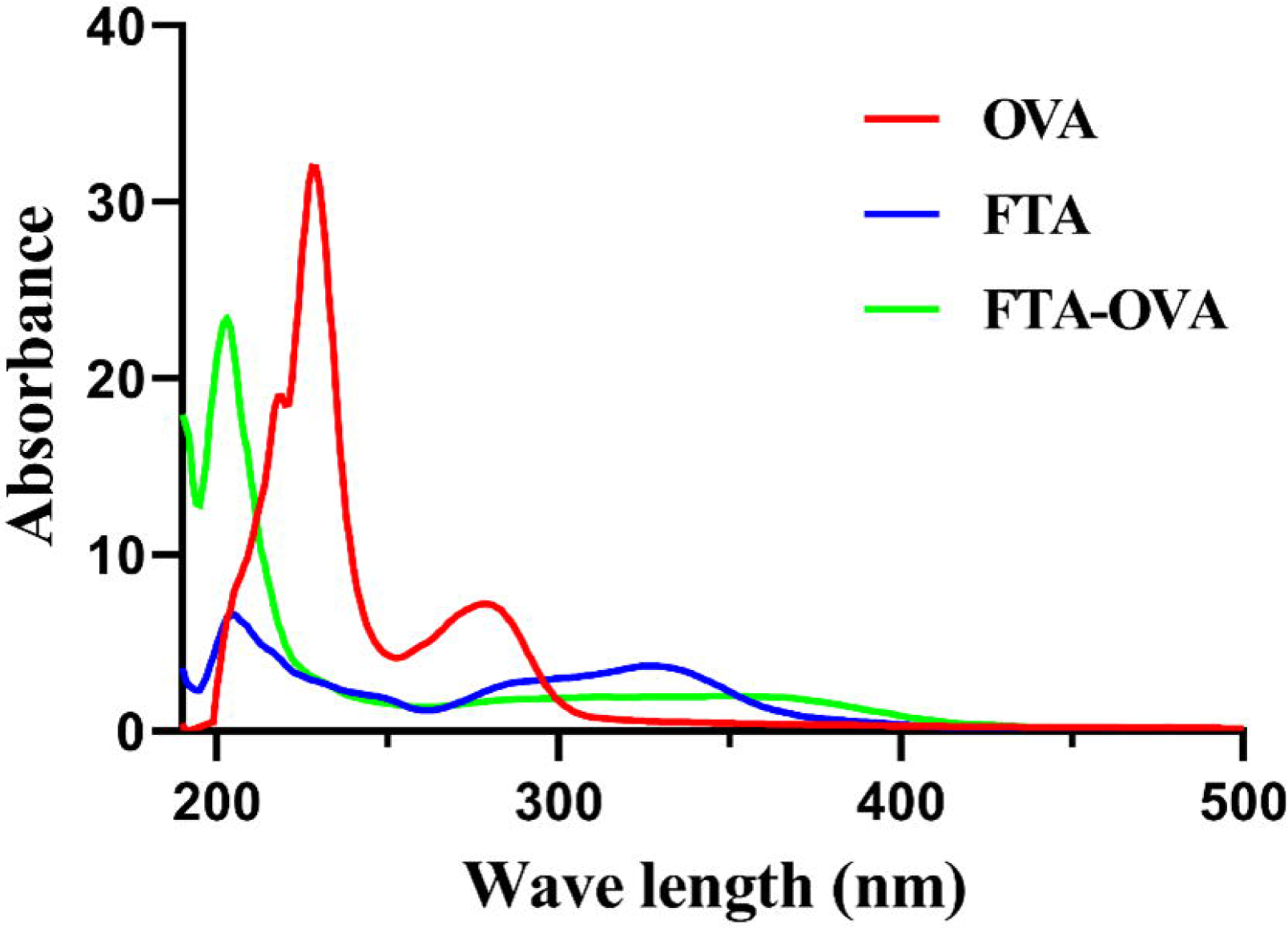
The artificial antigen FTA-OVA has been synthesized. Succinic anhydride method and carbodiimide method were used to synthesize FTA-OVA. The spectrum of three substances. Red indicated the spectrum of OVA, blue represented the spectrum of FTA and green was the spectrum of FTA-OVA.

### 3.5. Expression and purification of IBV S1 protein, S2 protein

HEK293T cells were transfected with plasmid encoding S1 protein or S2 protein, after 48 h incubation, the protein was collected and purified. The result of western blotting showed that there were approximately 100 kDa and 90 kDa bands indicated S1 protein and S2 protein, respectively (Fig.6A, Fig.6B). SDS-PAGE showed the result of purification. The lanes 1, 2 represented unpurified lysis solution and flow-through liquid, respectively (supplementary in Fig.S1A, Fig. S1B). In purification of S1 protein or S2 protein, lanes 3-4 (supplementary in Fig.S1A) or lanes 3-8 (supplementary in Fig.S1B) were collected after 100 mM imidazole elution, lanes 5-7 (supplementary in Fig.S1A) or lanes 9-12 (supplementary in Fig.S1B) were collected after 500 mM imidazole elution, S1 protein or S2 protein were mainly present in the eluent of lane 6 (supplementary in Fig.S1A) or lane 9 (supplementary in Fig.S1B).

**Fig.6.**
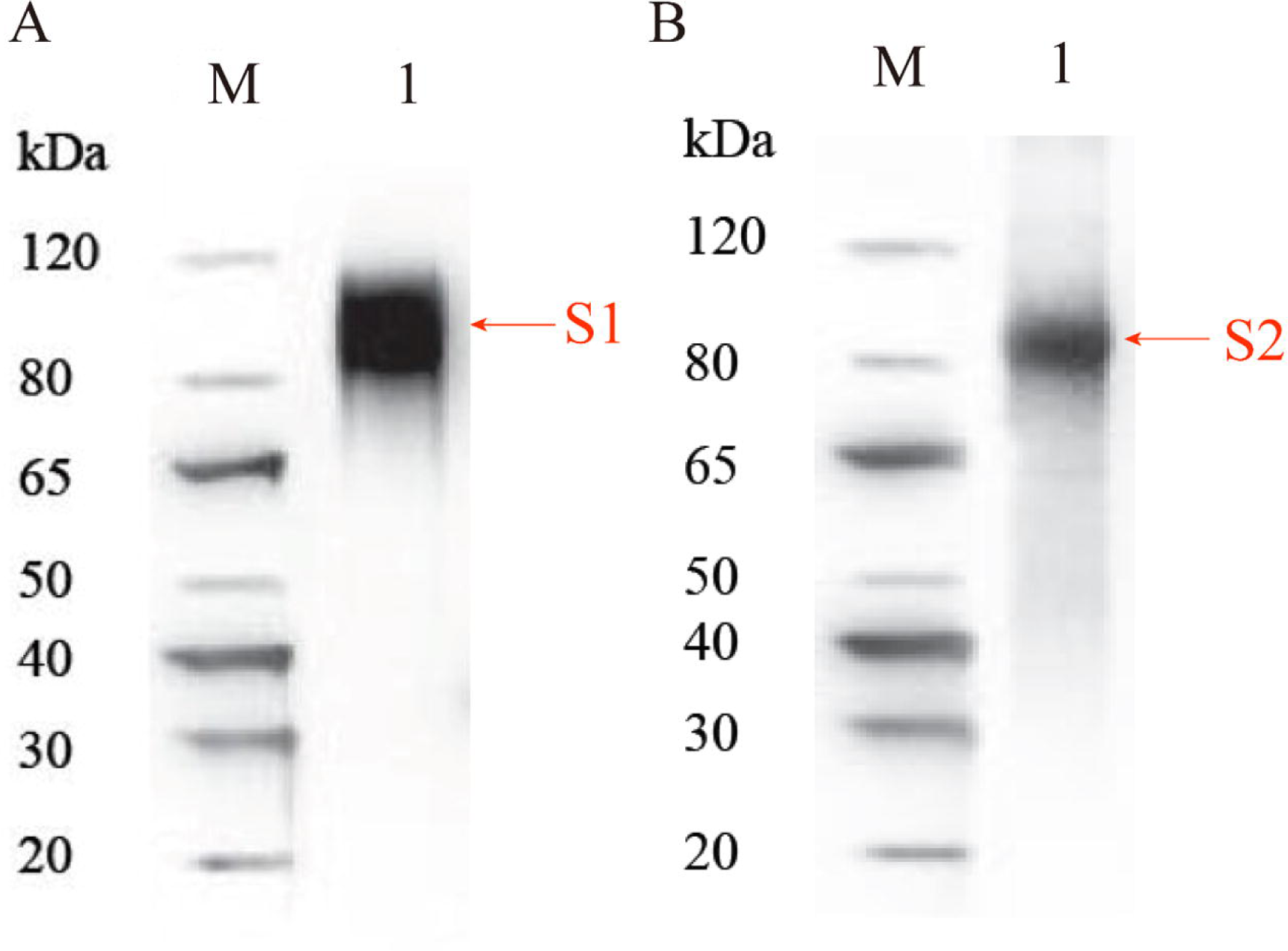
The expression and purification of S1 protein and S2 protein. (A) Western blotting showed that there was a band at approximately 100 kDa, indicated S1 protein. (B) Western blotting showed that there was a band at approximately 90 kDa, indicated S2 protein.

### 3.6. FTA bound to IBV S1 subunit but not S2 subunit

After the conjugation of FTA and OVA, FTA was used as a coating antigen for ELISA, then 800TCID_50_ IBV was added and ratio diluted 2-fold. DMEM contain 2% FBS was set as the control. The result of ELISA showed that the ratio of OD_450nm_ of positive/negative (P/N) increased following the amount of IBV added, revealed that FTA could indeed interact with IBV particles (Fig.7A). When the purified S1 protein was added in same method, the ratio of OD_450nm_ of P/N also increased (Fig.7B), preliminary identified that S1 subunit was bound to FTA.

**Fig.7.**
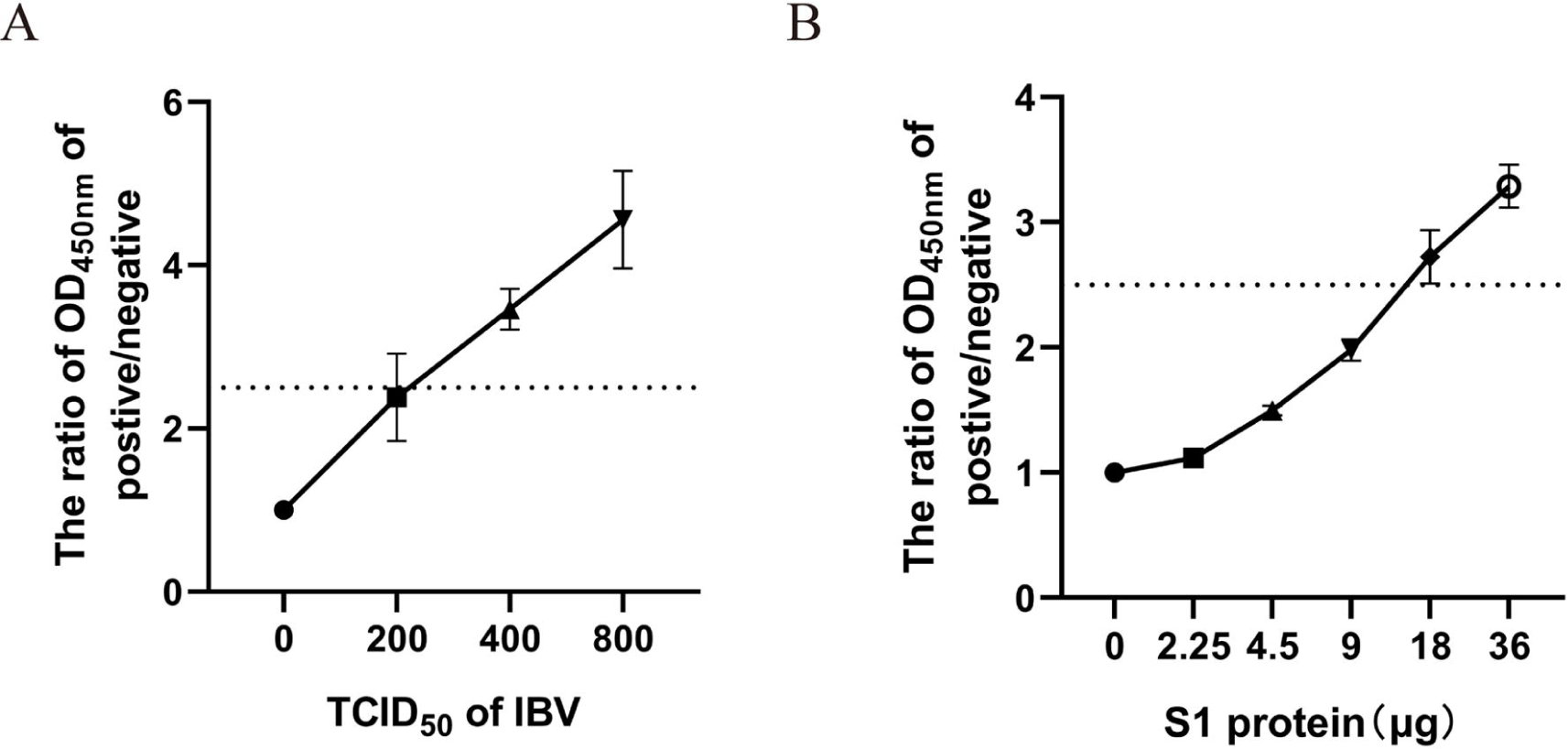
FTA interacted with IBV particles and S1 protein. FTA-OVA was coated on the ELISA plate as a determined concentration, then added (A) virus or (B) S1 protein with twice dilution ratios, the ratio of OD_450nm_ of positive/negative increased following the amount of virus or protein added. (A) The DMEM culture medium contain 2% FBS or (B) PBS was set as the control.

ITC was performed to further determine the interaction between FTA and S1 subunit. In the upper part of figure 8A, as the amount of FTA titration increased, the absorption heat of S1 and FTA reactions decreased, indicated that the binding between FTA and S1 protein tends to be saturated. In the lower part of figure 8A, the fitting curve was close to the “S” shape, and it was found that as the number of moles of FTA titration increased, the reaction between the two substances had a maximum rate, indicated that FTA interacted with the S1 subunit of IBV (Fig.8A).

**Fig.8.**
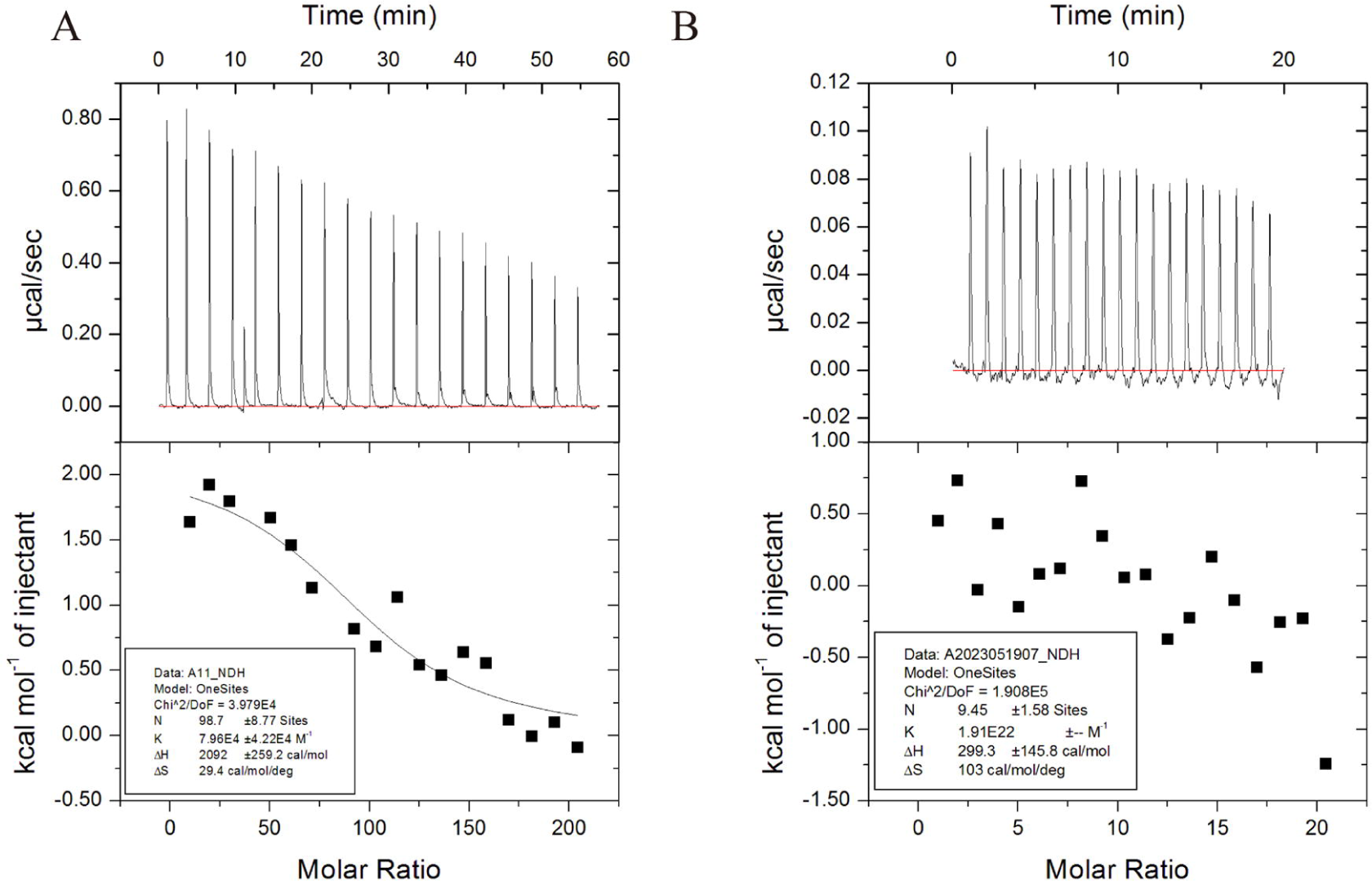
The results of FTA titrated S1 protein or S2 protein. (A) S1 protein was titrated by FTA. In the upper figure, the heat absorbed was decreased with adding FTA. In the lower figure, the fitting curve was close to the “S” shape, and there was a max gradient. (B) S2 protein was titrated by FTA. There is no observed regular peak, almost no heat change, and the scatter diagram was unordered.

There is a report found that IBV S2 subunit has a certain promoting effect on viral adsorption^[11]^. ELISA and ITC were performed and the results excluded the influence of S2 subunit on IBV attachment. The result of ELISA showed there were no significant changes of the ratio of OD_450nm_ of P/N when added S2 protein (data not shown), and when S2 protein was titrated with FTA, no regular peak was observed, almost no heat change, and the scatter diagram was unordered (Fig.8B), suggested FTA did not bind to IBV S2 subunit.

### 3.7. Alignment of FTA and IBV S protein

Using Autodock software, FTA might form hydrogen bonds with serine (aa 454) and tyrosine (aa 485) of IBV S protein (Fig.9A), and tyrosine (aa 434), threonine (aa 440), glycine (aa 441), serine (aa 469 and aa 471) aspartate (aa 473) interacted with FTA (Fig.9B). The donors and acceptors of hydrogen bond between FTA and IBV S protein were showed (Fig.9C). These amino acids all locate at S1 subunit of IBV S protein, and S1 subunit mediates IBV attachment, corresponded to the results of plaque assay and RT-qPCR previously.

**Fig.9.**
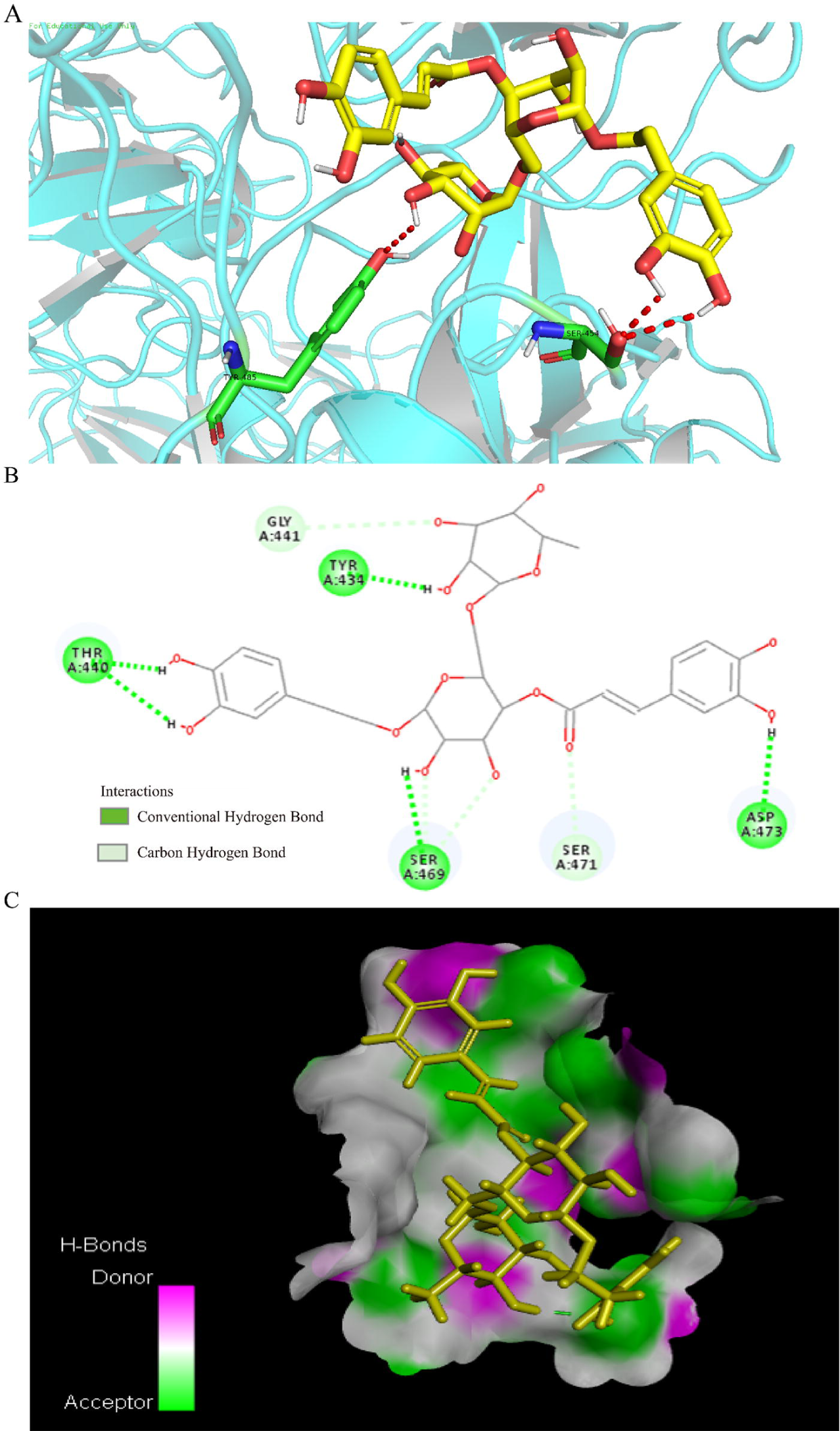
Docking results of FTA and IBV S protein. (A) FTA bound to serine (aa 454) and tyrosine (aa 485) of S protein by hydrogen bond (red dashed line). (B) Docking results showed that tyrosine (aa 434), threonine (aa 440), glycine (aa 441), serine (aa 469 and aa 471) and aspartate (aa 473) interacted with FTA. Dark green line indicated that FTA interacted with amino acids by conventional hydrogen bond, light green line indicated that FTA interacted with amino acids by carbon hydrogen bond. (C) The result of FTA bound to S protein showed the donors and acceptors of hydrogen bond.

**Fig.10.**
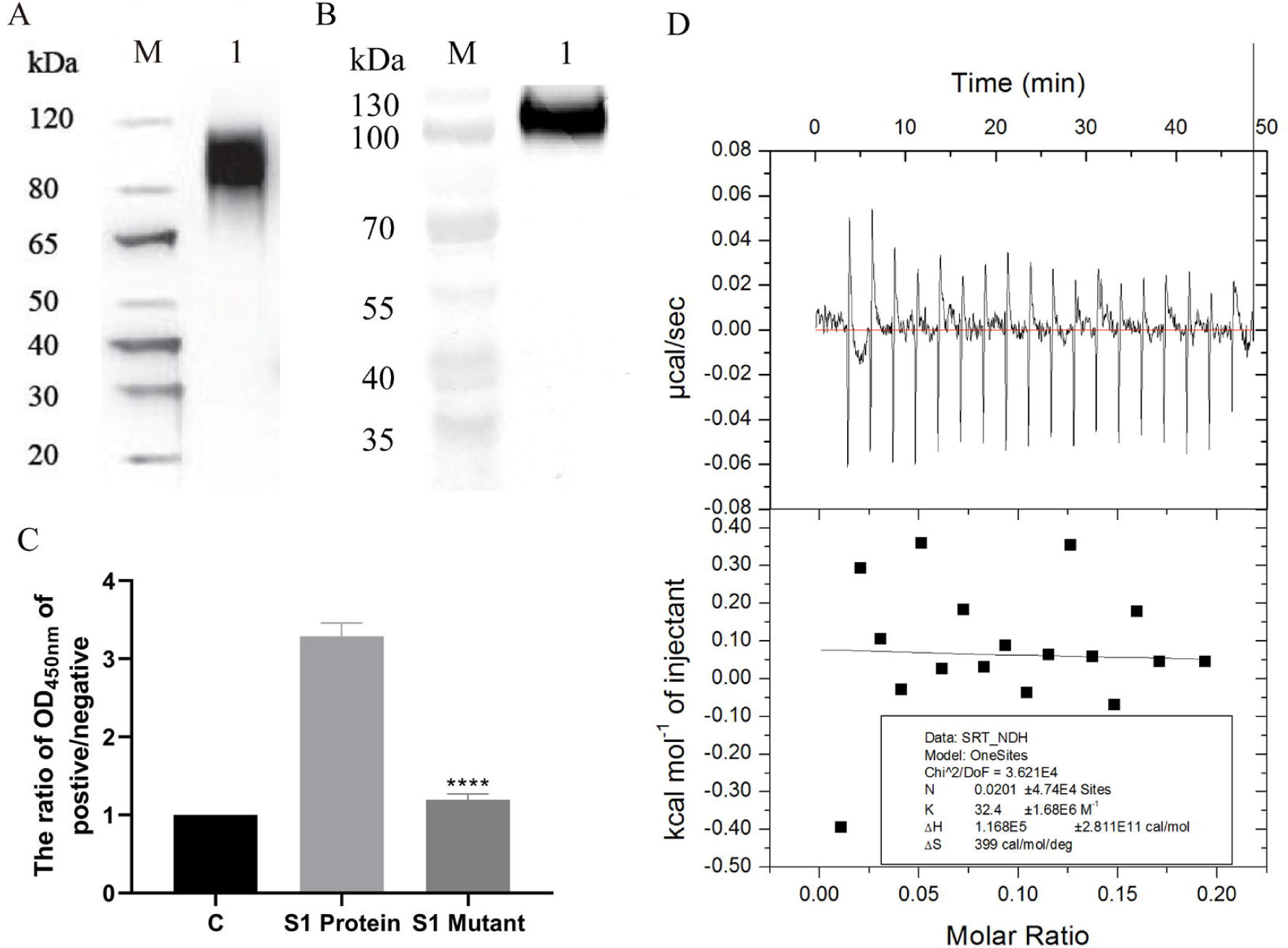
The expression of S1 mutant was showed by western blotting and the comparison of affinity of FTA bound to S1 protein or S1 mutant. (A, B) After the amino acids of S1 protein were changed into another amino acids with similar structure and properties, the molecular weight of S1 mutant and S1 protein was close, approximately 100 kDa. (C) The ratio of OD450nm of P/N was decreased when S1 protein was mutated. ****p < 0.001, compared with S1 protein. (D) S1 mutant was titrated by FTA. Unable to determine whether it is an endothermic or exothermic reaction, and there was almost no heat change.

### 3.8. IBV Beaudette S1 mutant loss binding ability with FTA

To explore the functions of these sites on FTA binding, S1 mutant was generated by mutating the eight amino acids into phenylalanine (aa 434), alanine (aa 440), alanine (aa 441), valine (aa 454, aa 469 and aa 471), asparagine (aa 473) and phenylalanine (aa 485) (supplementary in Table S1). Firstly, the expression of S1 mutant was detected by western blotting. Corresponded to the S1 protein, S1 mutant was approximately 100 kDa (Fig.10A, Fig.10B). Then the interaction between S1 mutant and FTA was explored by ELISA and ITC. When the molar concentration of S1 protein and S1 mutant was close, the ratio of OD_450nm_ of P/N of S1 mutant was decreased. What’s more, the result of ITC showed that there was almost no heat change when FTA and S1 mutant titration (Fig.10C, Fig.10D), indicated that the affinity disappeared after mutated those sites, revealed that FTA could bind these sites, but not found which one was FTA exactly bound.

## 4. Discussion

Coronavirus infect cells can be roughly divided into the following steps: attachment, entry, replication, assembly, and release^[12]^. To explore the anti-viral mechanism of FTA, plaque assay and RT-qPCR were performed to test the effect of FTA on each step of IBV infection. It was found that FTA inhibited IBV attachment to surface of Vero cells membrane. Pretreated cells with FTA, IBV infection couldn’t be prevented; but once IB virus incubated with FTA, the infection rate decreased. These results indicate that the inhibition of IBV attachment by FTA is due to interaction with viral particles, rather than with cellular receptors. Spike protein is the largest structural protein of coronavirus, which forms rod like protrusions on the surface of viral envelope and exists in the form of dimer or trimer^[4]^. The S protein of IBV is a highly glycosylated Class I fusion protein^[13]^. It has a special pentapeptide motif RRFRR on its amino acid sequence, which can be recognized and cleaved by the furin-like protease in the host cell. Subsequently, the S protein is divided into S1 subunit containing approximately 500 amino acids and S2 subunit containing approximately 600 amino acids^[10, 14]^. S1 subunit is an important part for IBV to exert adsorption function, and all known receptor binding domains (RBDs) are located on the S1 subunit^[15]^. According to the results above, S1 subunit might be the binding part of FTA. The docking results showed that there were eight sites where FTA probably bind to S protein, and the sites were all located at S1 subunit. S2 subunit contains an extracellular domain, a transmembranedomain, and an intramembrane domain, which are anchored to the viral envelope through the intramembrane domain to mediate the fusion of the viral envelope and cell membrane^[16]^. What’s more, S2 subunit could promote virus attachment^[11]^. Then S1 and S2 protein were expressed and purified, and molecular interaction assays were performed. Both ELISA and ITC results indicated that FTA interact with S1 subunit rather than S2 subunit. In order to further confirm that FTA bound to the eight amino acids, we chose other amino acids with similar structures to replace the amino acids of wild S1 subunit. As table S1 showed, the R group of amino acids of S1 mutant were different from amino acids of S1 wild, and these mutated amino acids might cause inability to generate hydrogen bonds or weakening of bond energy with FTA. Such as tyrosine, which was converted to phenylalanine, lost phenolic hydroxyl groups, might interfere with the formation of hydrogen bonds, might result in S1 mutant loss affinity to FTA, but which amino acid plays a decisive role is still unknown.

When these sites were mutated into another amino acids with similar property, the affinity of mutant to FTA was lower than S1 protein (in similar weight), so we considered that FTA might block the interaction between S protein and certain receptors, rather than known receptors. The effect of each site which we found in this study should be explored in the future.

## 5. Conclusion

The molecular mechanism of anti-IBV infection and the targets of FTA were identified. FTA prevents IBV infection by blocking viral attachment and contacts with the S1 subunit of IBV, but not with the S2. At last, the bind sites were identified by comparing the affinity of wild S1 and S1 mutant.

## Declarations

## Ethics approval

Ethical approval is not required.

